# Expression QTLs in single-cell sequencing data

**DOI:** 10.1101/2022.08.14.503915

**Authors:** Ariel DH Gewirtz, F William Townes, Barbara E Engelhardt

## Abstract

Single nucleotide polymorphisms (SNPs) are important drivers of gene expression variation and downstream phenotypes including disease risk. Single-cell RNA-sequencing (scRNA-seq) allows an unprecedented exploration of cell-type specific associations between gene expression levels and genotypes, but current methods rely on pseudobulk approaches that use composite expression values across cells and often use summary statistics within cell types, ignoring information across cell types and assuming cell type labels are accurate. Here, we extend our method, telescoping bimodal latent Dirichlet allocation (TBLDA), that identifies covarying genotypes and gene expression values when the matching from samples to cells is not one-to-one in order to allow cell-type label agnostic discovery of eQTLs in noncomposite scRNA-seq data. In particular, we add GPU-compatibility, sparse priors, and amortization to enable fast inference on large-scale scRNA-seq data. We apply single-cell TBLDA (scTBLDA) to 400K cells from 119 individuals with systemic lupus erythematosus and examine properties of features from each modality across the estimated latent factors. We use linked genes and SNPs to identify 205 cis-eQTLS, 66 trans-eQTLs, and 53 cell type proportion QTLs, which we then compare against prior studies of immune-cell eQTLs. Our results demonstrate the ability of scTBLDA to identify genes involved in cell-type specific regulatory processes associated with SNPs in single-cell data.

## 1 Introduction

Single nucleotide polymorphisms (SNPs) are known to regulate gene transcription and are important drivers of population-level variation in gene expression. Until recently, most expression quantitative trait loci (eQTLs) were mapped using gene expression data from bulk RNA-sequencing data. Bulk RNA-seq quantifies average expression levels over an often heterogeneous pool of cell types in each sample, which misses associations exclusively in rare cell types and confounds associations by cell-type heterogeneity through differential gene expression across cell types [1]. Single-cell RNA-seq (scRNA-seq) has revolutionized the granularity at which we can explore expression patterns and transcriptional regulation by examining expression within individual cells instead of across a pool of heterogeneous cells. Specifically, scRNA-seq provides a fine-grained view into cell-type specific variation in gene expression, which is impossible to characterize accurately from bulk data.

However, scRNA-seq requires careful statistical modeling given its sparsity and increasingly large sample size. Moreover, there are multiple gene expression observations (typically hundreds to thousands of cells) for every genotype profile in an eQTL study. Many eQTL mapping models require Gaussian-distributed expression data, including the most common linear regression-based model. Common normalization procedures for scRNA-seq data, such as log-transformation, have been shown to distort the count data and lead to spurious differential expression and gene variability [2, 3].

Generally, eQTLs are mapped by testing for linear association between a single SNP and a single gene across a matched number of samples, with each individual contributing one SNP and one gene expression measurement. In single-cell sequencing data, this is accomplished by summarizing gene expression levels from a collection of cells using the sum of each gene’s expression across those cells, an approach referred to as *pseudobulk.* Pseudobulk approaches do not take advantage of the rich variation in expression across and within cell types and samples. These simplistic one-to-one mappings also ignore the larger, deeply intricate regulatory networks that drive the associations, may be overly reliant on cell type annotations, and may impose relationships such as correlation across expression. Alternatively, because of the large proportion of zeros in scRNA-seq, if the one-to-one association methods added a ‘pseudo’ sample for every cell measured, all paired to the same SNP value for the respective donor, the proportion of zeros in the gene expression measurements would overwhelm association signals and few, if any, associations would be found.

One approach to integrate multiple genes is to cluster them into co-expressed modules and assume that each cell’s expression vector is made up of a linear combination of latent expression programs. Several models have been built for single-cell data that identify latent gene expression or epigenetic factors and associate them with important biological states such as cell type and differentiation progression [4, 5, 6, 7, 8]. Although not implemented in the aforementioned methods, those latent factors could then be tested for association with a SNP in a one-to-one manner in order to detect pleiotropic genetic effects. To account for correlation among SNPs due to linkage disequilibrium (LD), some approaches have also considered composite SNP measures or including multiple SNPs in tests for association [9].

Because of these limitations, previous work to map eQTLs using expression data from scRNA-seq generally requires data pre-normalization and uses pseudobulk methods, has been limited to cis-eQTLs, relies on cell-type labels and subsets of cells in a sample to quantify gene expression levels, and does not jointly model genotype and expression data beyond the one-to-one association tests [10, 11, 12, 13, 14, 15, 16].

We include all of these ideas in our probabilistic framework telescoping bimodal latent Dirichlet analysis (TBLDA), using both data modalities to jointly determine a set of latent factors that each correspond to a collection of covarying genes and SNPs. In prior work on TBLDA, the estimated factors contain sets of associated genes from multiple tissues using bulk RNA-seq (instead of many cells from the same individual) and SNPs measured in a single donor [17]. In this paper, we expand the TBLDA framework to enable efficient inference across large-scale scRNA-seq data sets. Our improved method, single-cell TBLDA (scTBLDA), allows read counts from each cell to be considered without aggregation and does not rely on subsetting the cells by potentially inaccurate pre-defined cell-type labels to create pseudobulk statistics. Furthermore, we add to TBLDA GPU-compatibility, sparse priors, and amortization during inference to enable fast application to population-scale scRNA-seq data.

We apply scTBLDA to a dataset of approximately 400,000 peripheral blood mononuclear cells (PBMCs) from individuals with systemic lupus erythematosus (SLE) [18]. We use the estimate scTBLDA model to identify and explore cell-type associated factors, including identifying unlabeled functional subtypes of two separate cell types in PBMCs. Next, we use linked genes and SNPs to map cis- and trans-eQTLs both within and across cell types in addition to mapping cell-type proportion QTLs. We validate these cis- and trans-eQTLs in existing PBMC study data, and further show evidence of cell-type specific eQTLs identified using our approach that were missed using bulk and pseudobulk methods.

## Methods

To enable adaptation to single-cell sequencing data, we developed three functionally-important modifications to the TBLDA model presented in prior work [17]: GPU-compatibility, a sparsity-inducing prior for the loading matrices, and amortization during inference. We refer to the new model as single-cell TBLDA (scTBLDA).

### Model Improvements

To facilitate realistic computational run times on ever-growing data sets, we extended our model to run on GPUs. With this extension, inference to fit our model on approximately 2,500 genes across 400,000 cells and 12,000 SNPs across 120 individuals using 50 latent factors takes around 30 minutes. We contrast this runtime with the original approach, which took approximately 24 hours using the same data.

Latent Dirichlet allocation (LDA) results are influenced by the per-factor feature hyperparameter; large values will encourage many features to define a factor while values < 1 encourage factors to be defined by a small number of features. We explicitly set all of these hyperparameters to symmetric values of 1/K, where *K* is the number of factors, to encourage sparsity in our estimated loadings. Increasing sparsity means that fewer features (SNPs, genes) are active per factor, yielding factors that are easier to interpret overall.

In the original model formulation, we use variational inference to estimate the factor proportions in each sample. Although feasible for medium-sized data, this clearly scales poorly with increasing sample size *L.* In our prior work, *L* = 5, 781 was the number of bulk RNA-seq samples. In order to accommodate single-cell data sets with L in the hundreds of thousands to millions, as we have here, we use amortization to vastly reduce the number of model parameters. Instead of learning a local variational approximation for every cell’s factor proportions *φ_ℓ_, ℓ* = 1,…, *L*, we learn a flexible function *ϕ_ℓ_* ≈ *f*(x_*ℓ*_) to estimate the parameter for each cell given its expression vector x_*ℓ*_ (Extended Methods). Specifically, we train an encoder with two fully connected hidden layers of 300 and 150 neurons, respectively. The model results are robust to changes in hidden layer sizes, learning rates, and activation functions (Tables S1, S2, S3). We use batch normalization to avoid component collapse, a common pitfall where many (or all) factors collapse into a single common mode [19]. Although other factor models had positive outcomes using both batch normalization and dropout [19], our model did not overfit to the training data—adding varying proportions of dropout only increased the test set loss and posterior expression root mean squared error (RMSE; Table S4).

### Top Feature Selection

Due to the encouraged sparsity in the posterior, we modified how we select the top features in each factor from the previous 2-Wasserstein-based metric [17]. To correct for the correlation between feature levels and factor weights, we conducted quantile regression on the factor loadings to adjust for allele frequency and average expression level (Fig. S1). This prevents features with higher counts from dominating the results, as all feature levels contribute an approximately equal proportion to the set of top features per factor (Fig. S2).

## Results

### Factor Associations with Covariates

We first identified the factors that are associated with known covariates *cell type* and *batch* (Fig 1). Although these covariates both drive gene expression variation within the data set, batch effects induce technical variation that we would like to separate out from biologically-driven signal. On average, 3.5 out of 50 factors per scTBLDA run were associated with batch and subsequently removed from downstream analysis. Overall, as we expect, factors were more associated with cell-type proportions than batch (Fig. S3). Out of 1320 cell-type associated factors, the majority were associated with CD4 T cells and CD14+ monocytes – 478 and 466, respectively. Megakaryocytes and dendritic cells had the fewest number of associated factors with 22 each. The number of associated factors per cell type was strongly correlated with the number of cells (Kendall’s rank correlation *τ* = 0.84; *p* ≤ 0.005). This suggests that, while analyses for rare cell types are generally underpowered, they may be improved by sharing strength across cell types.

**Figure 1:**
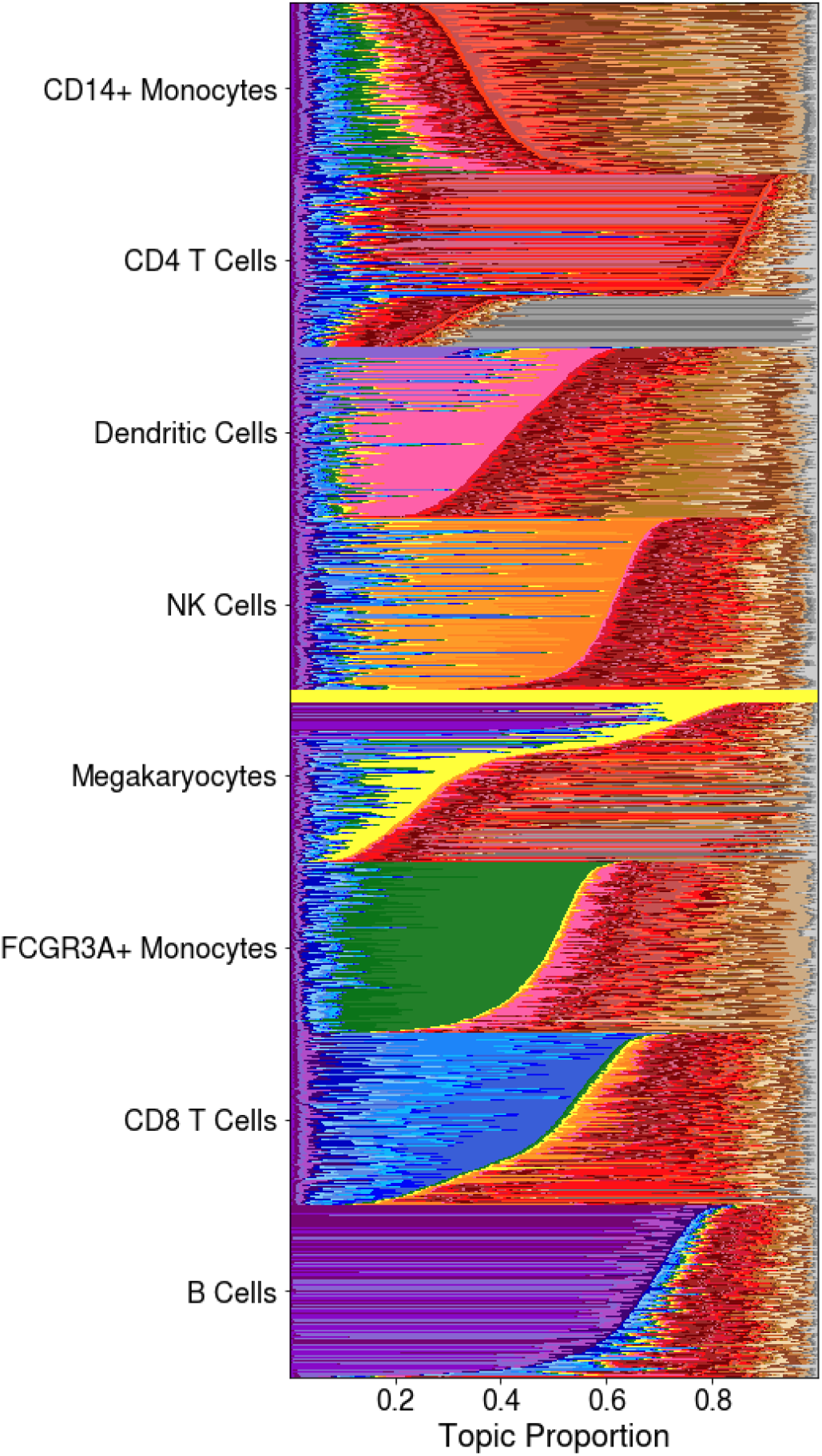
Visualization of Φ (per-cell factor proportions) for 1000 randomly-chosen cells from each cell type from the scTBLDA run using chromosome 12 SNPs. Cell-type specific factors are colored in shades of purple, blue, green, yellow, orange, pink, red, and brown, while factors associated with batch are colored gray. Cells are first ordered by the provided cell-type label and then within each cell type by decreasing proportion of cell-type associated factors. This run is representative of scTBLDA runs using SNPs on other chromosomes.

### scTBLDA Factors Capture Cell Subtypes

Just as bulk tissue samples encompass a heterogeneous collection of cells, cell types are themselves umbrella terms, clustering largely-similar cells that can still be separated into distinct cell subtypes. While megakaryocyte-labeled cells were the second rarest, composing just under two percent of all cells in this sample, they had striking intra-cell type factor variation (Fig 1). For example, we identified four megakaryocyte cell sub-populations: one entirely described by the megakaryocyte factor, B cell-like, CD8 T cell-like, and CD4 T cell-like (Fig 2A). Loadings for the three most variable factors across megakaryocytes separated the cells into three distinct clusters (Fig 2B). The most informative genes in each factor mirror the cell-type associations. In the B cell-like subgroup, the most informative gene was *MS4A1,* which encodes a B-lymphocyte surface antigen. The top gene in the full megakaryocyte factor, *PPBP,* encodes a CXC cytokine growth factor. Correspondingly, the second most informative gene in both the CD4 and CD8 T cell-like subpopulations was *P1K3IP1,* which inhibits T cell activation. While it is possible that some megakaryocytes were mislabeled, a notable proportion of each cell’s topic vector *ϕ_ℓ_* is assigned to the megakaryocyte-associated factor; this stands in stark contrast to, for example, the B cell topic vectors (Fig 2A).

**Figure 2:**
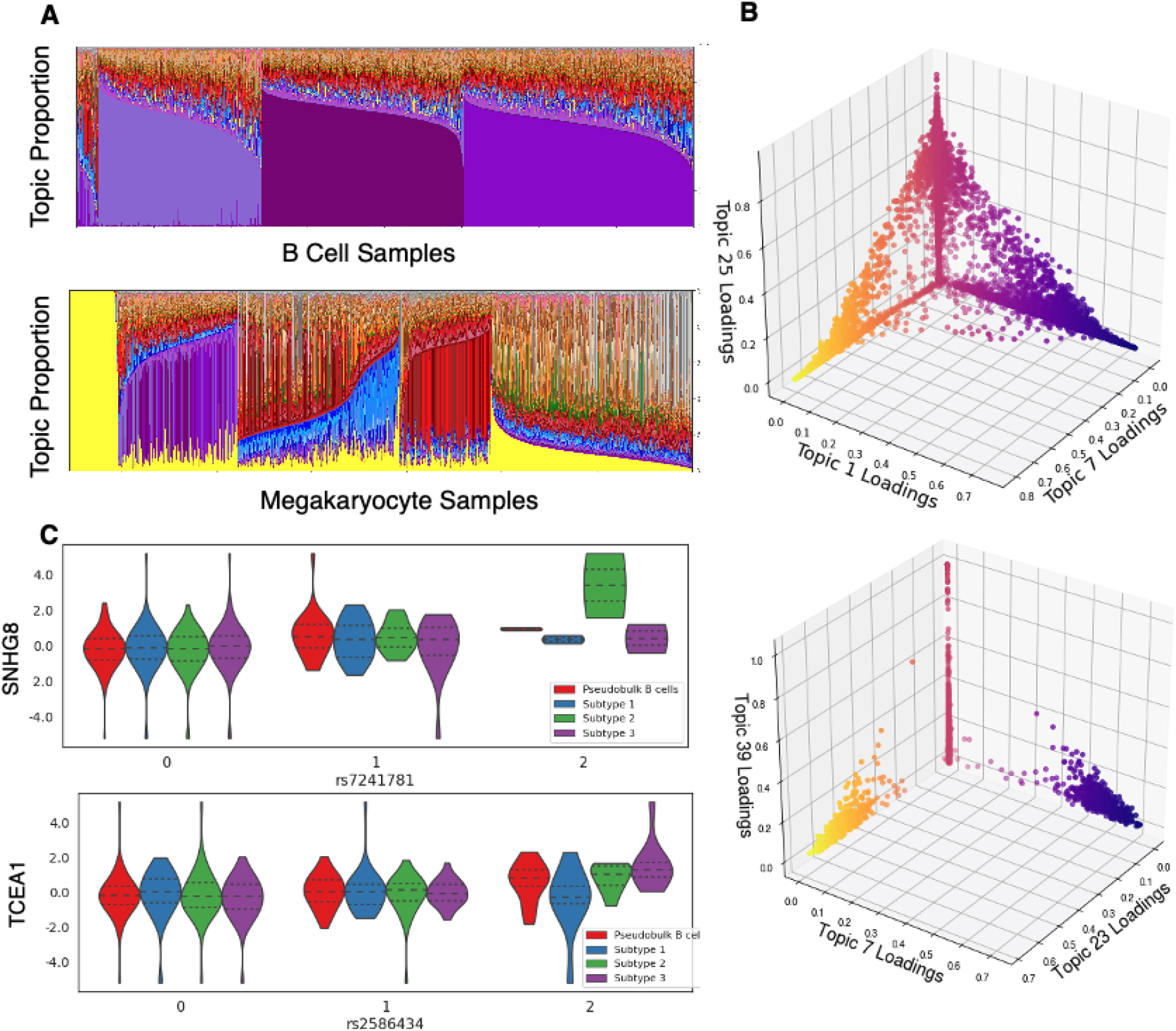
Cell subtype expression variation and associations with genetic variants. **A.** Visualization of Φ (per-cell factor proportions) for 8000 randomly-chosen B cells and all megakaryocytes from the scTBLDA run using chromosome 12 SNPs. Topics are colored the same as in Fig 1. Cells are ordered by dominant topics. **B.** 3-D visualization of B cells (top) and megakaryocytes (bottom) determined by their expected factor loadings for the three most variable factors within each cell type. Topics 1, 7, and 25 are associated with B cells, topic 39 is associated with megakaryocytes, and topic 23 is associated with CD4 T cells. **C.** Example trans-eQTLs that are specific to a subset of B cells, including the association between *SNHG8* and rs7241781 (p ≤ 2.1 × 10^-5^) and *TCEA1,* which is involved in DNA binding, and rs2586434 (p ≤ 1.7 × 10^-7^). The x-axis represents the number of copies of the minor allele at each locus.

Next, we explored intra-cell-type factor variation among B cells. We identified three B cell sub-populations, each characterized by a unique B cell-associated factor containing a majority of the cell’s weight (Fig 2A). In contrast to the megakaryocytes, the B cell loadings form a continuum across the three most variable factors (Fig 2B). This may be because the three B cell subtypes share 73% of their most informative genes. However, only 7% of the top SNPs are shared across these factors, suggesting potentially different regulatory mechanisms across overlapping gene sets.

To test whether these three B cell factors captured subtype-specific regulatory associations, we mapped eQTLs separately in each cell subtype using the top genes and SNPs for each factor on held-out data (see Extended Methods). Out of 28,266,063 tests, we found 19 trans-eQTLs comprising 16 unique trans-eGenes and 19 unique eVariants that were only present in one of the cell subtypes (BH FDR ≤ 0.2); three eQTLs include the lncRNA *SNHG8* (Fig 2C). Through this analysis, we not only emphasize the importance of context even within scRNA-seq data from a single cell type but also illustrate how scTBLDA is particularly well-suited for this type of exploration.

### Functional Characterization of Cell-Type Associated Features

We compiled sets of genes that were repeatedly top-ranked across runs in factors associated with each cell type (Fig 3A; see Extended Methods). In line with cell-type abundance, CD14+ monocytes and CD4 T cells had the largest gene sets (1405 and 949, respectively) while the dendritic cell gene set was the smallest (160 genes). All cell-type associated gene sets had immune-relevant enrichments using KEGG, Reactome, and GO biological process gene sets (one-sided hypergeometric test, BH FDR < 0.1; Table S5). For example, the most significant enrichment for natural killer cell genes was the KEGG natural killer cell mediated cytotoxicity gene set (BH FDR ≤ 2.5 × 10^-4^) and the second largest enrichment for the most informative megakaryocyte-associated genes was the Reactome platelet activation signaling and aggregation pathway (BH FDR ≤ 1.0 × 10^-6^). Several cell-type associated gene sets also had regulatory gene set enrichments relating to transcription, translation, and splicing. Among others, the B cell, CD8 T cell, and CD4 T cell sets were all enriched for Reactome 3 UTR mediated translational regulation and Reactome nonsense-mediated decay enhanced by the exon junction complex genes (BH FDR < 1 × 10^-16^), and the CD14+ monocyte set was enriched for GO formation of translation pre-initiation complex genes (BH FDR ≤ 0.09).

**Figure 3:**
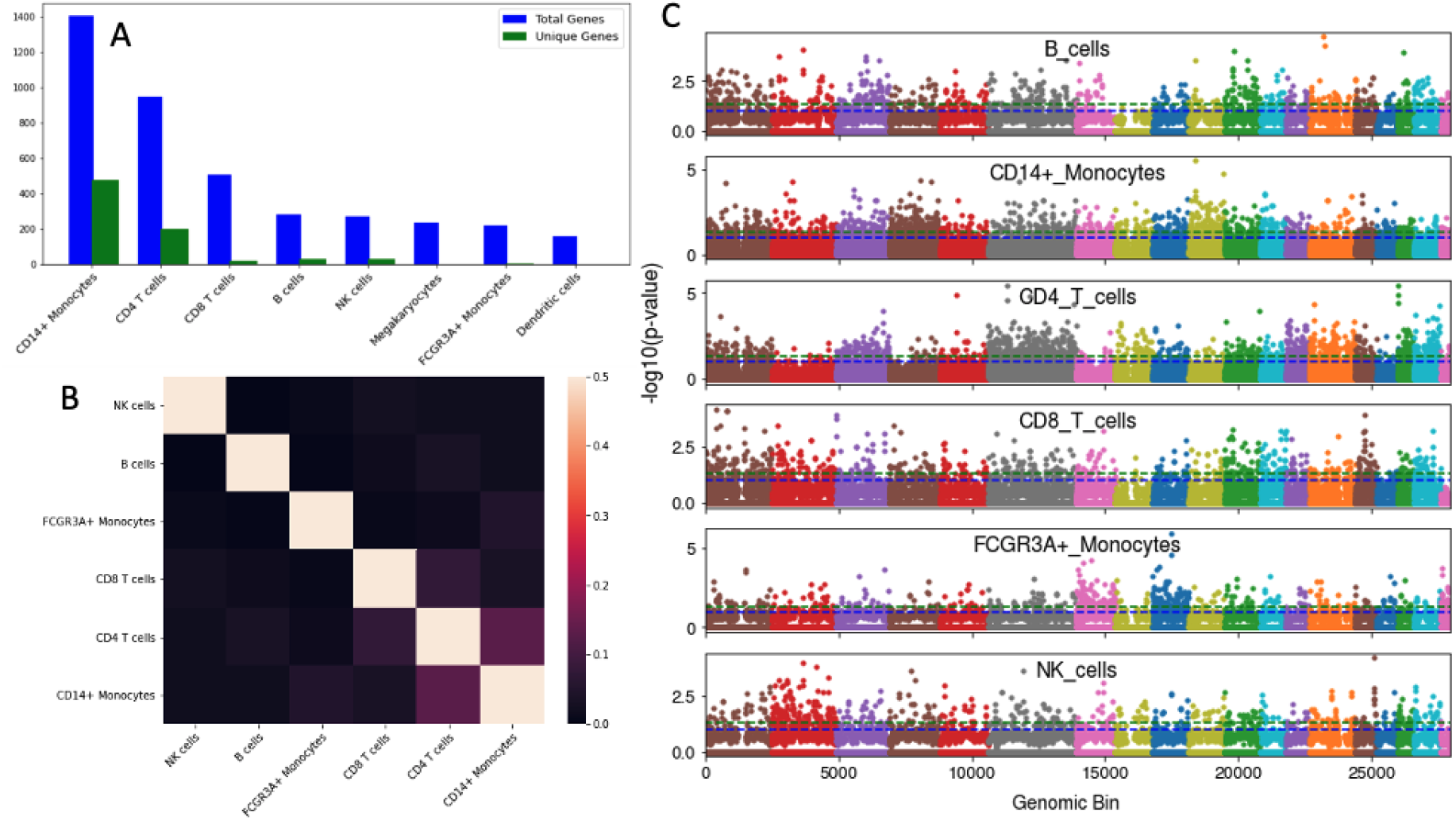
Cell-type associated feature analysis. **A.** Bar plot depicting the number of total and unique genes associated with each cell type. **B.** Jaccard similarity measure between the cell-type associated SNP sets. CD4 and CD8 T cells have overlapping SNPs as well as CD14+ and FCGR3A+ Monocytes. **C.** Genomic bin enrichment of cell-type associated variants. Bins are colored by chromosome; the green and blue dashed lines correspond to significance levels of 0.05 and 0.1, respectively.

Beyond composite cell-type associated gene sets, the top genes identified within individual scTBLDA factors were also enriched across GO, KEGG, and Reactome gene sets. Notably, 22 individual megakaryocyte-associated factors were all enriched in the Reactome RNA Pol I promoter opening genes (BH FDR ≤ 0.1). Finding such enrichment among intra-factor top-ranked provides an outside line of evidence that these genes may be co-regulated.

Next, we explored SNPs that were frequently top-ranked in cell-type associated factors and tested for enrichment in 35 functional annotation classes [20] (see Extended Methods). All cell types except for megakaryocytes and dendritic cells had cell-type associated SNP sets (Fig 3B). The CD4 T cell, CD14+ Monocyte, FCGR3A+ Monocyte, and B cell SNPs were all enriched in enhancer-associated markers, while NK cell SNPs were enriched for transcription start sites and DNAse hypersensitive sites (one-sided Fisher’s exact test, *p ≤* 0.04). The union of all cell-type associated SNPs was enriched for 13 functional annotations including super enhancers and enhancer-associated H3K4me1 (one-sided Fisher’s exact test BH FDR ≤ 0.004, Table S6). This suggests that scTBLDA identifies top-ranked SNPs enriched in enhancer-associated regions that tend to colocalize with trans-eQTLs, echoing previous findings [17, 1].

To test whether the cell-type SNP sets clustered together in particular genomic regions, we looked for enrichment in genomic bins of size 250 Kb within each chromosome using a sliding window of 100 Kb. All cell-type associated SNP sets were enriched in at least one bin (one-sided Fisher’s exact test, BH FDR < 0.1; Fig 3C, Table S7). Such bin enrichments suggest potential regulatory hotspots, DNA regions that affect the expression levels of many genes [21]. Notably, many of the cell-type associated SNP sets were enriched in consecutive bins, indicating wider genomic regions potentially active in gene regulation for specific cell types. FCGR3A+ monocyte SNPs were enriched in two consecutive bins starting at chromosome 10 position 66665030 (BH FDR ≤ 0.02), B cell SNPs in two bins beginning at chromosome 15 position 74621207 (BH FDR ≤ 0.022), CD4 T cell SNPs in two bins starting at chromosome six position 68253012 (BH FDR ≤ 0.014), chromosome 19 position 1589282 (BH FDR ≤ 0.004), and chromosome 21 position 42306585 (BH FDR ≤ 0.08), CD8 T cell SNPs at chromosome one position 79391536 (BH FDR ≤ 0.066), and CD14+ Monocyte SNPs at chromosome 11 position 24892155 (BH FDR ≤ 0.075). The largest consecutive region enrichment belonged to the CD4 T cell SNPs—enriched in three consecutive bins starting at chromosome 21 position 17106585 (BH FDR ≤ 0.095). Together, these results demonstrate that scTBLDA is able to identify broad genomic regions that contain many SNPs with cell-type specific associations.

### Exploring the Relationship between Linked Genes and SNPs

First, we tested for positional overlap among the the most informative features in each estimated factor. Four out of 22 runs had factors for which the top genes were enriched for genes on the same chromosome as the genotype data (Fisher’s exact test BH < 0.2; chromosomes 6, 12, 13, 19). Three runs had marginal enrichment among the union of top genes across all factors for origin on the same chromosome as the genotype data (one-sided Fisher’s exact test, p-value *p ≤* 0.2; chromosomes 14, 19, 21). Taken together, these indicate that the majority of scTBLDA factors capture trans relationships—those between genes and SNPs on different chromosomes [1].

We further explored the relationship between SNPs and genes loaded onto common factors by mapping eQTLs (see Extended Methods). We mapped associations between the top SNPs and genes only in factors that were associated with a particular cell type. In total, we found 205 cis-eQTLs including 90 unique cis-eGenes and 169 unique cis-eVariants, and 42 intra-chromosomal long-range cis-eQTLs including 39 unique cis-eGenes and 42 unique cis-eVariants (Bonferroni FDR < 0.2; Table S8). We also found 66 trans-eQTLs including 48 unique trans-eGenes and 65 unique trans-eVariants (Fig 4; Table S8). Although the cis-eQTLs mostly contained protein-coding eGenes (96%), the trans-eQTLs were almost evenly split with lincRNA eGenes (53% to 47%). This composition of protein-coding and linkRNA within the cis- and trans-eQTLs agrees with our previous TBLDA findings using GTEx data [17]. Of the cell types, CD14+ monocytes had the most cis-eQTLs (67), dendritic cells had the most long-range cis-eQTLs (10), and CD8 T cells and NK cells tied for the largest number of trans-eQTLs (11 each). To ensure that the signal in scTBLDA shared components is meaningful, we tested all associations between top features while permuting expression and covariate values. We found enrichment across all associations (Fig S4; Fig S6).

**Figure 4:**
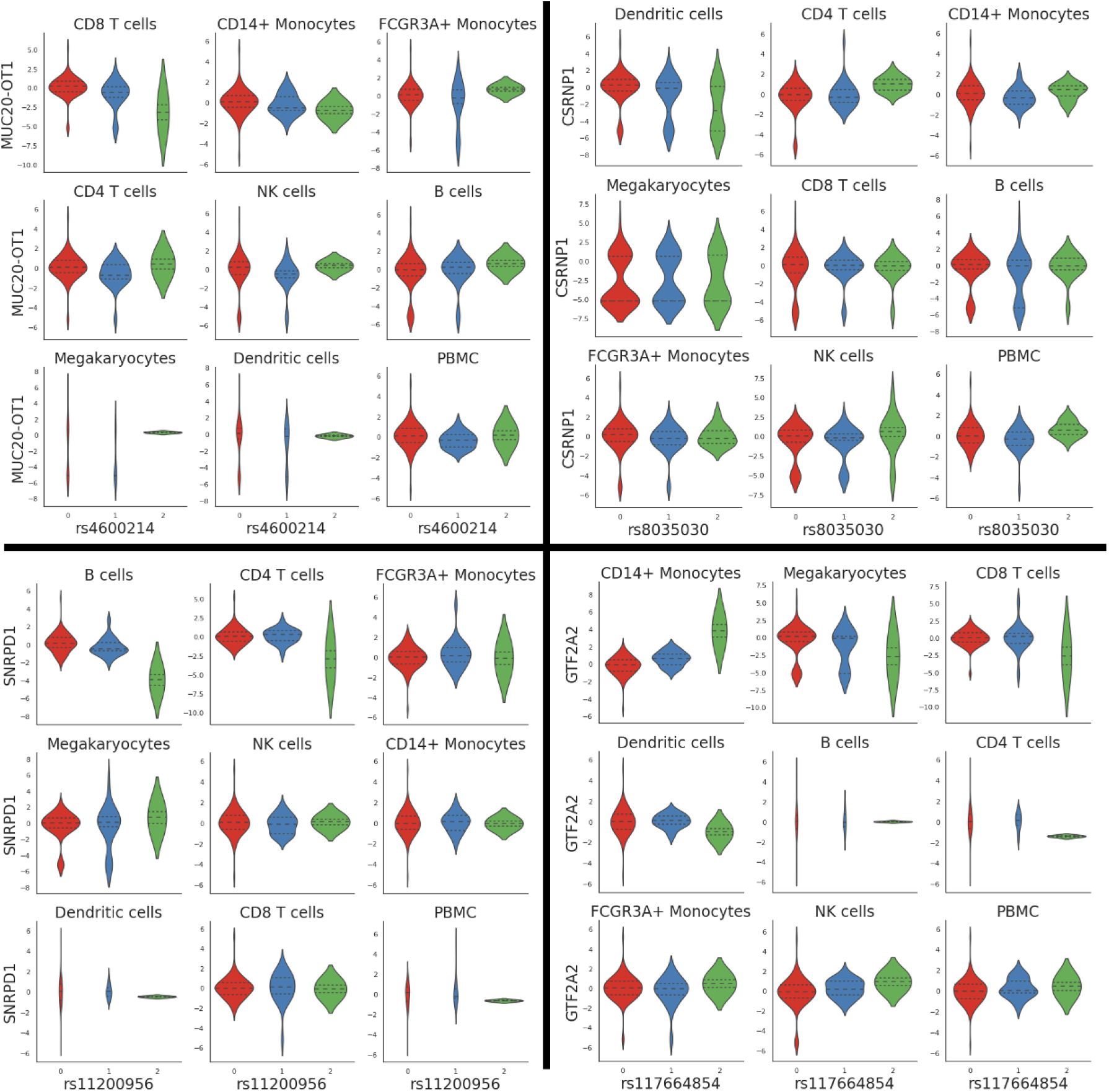
Example trans-eQTLs in each cell type and pseudobulk PBMCs. The y-axis represents quantile-normalized gene expression values and the x-axis is the number of copies of the minor allele at the labeled variant. Each point represents a single cell. We find cell-type specific trans-eQTLs that are masked in pseudobulk PBMCs. We also find opposite directional effects from a given allele in different cell types (as in associations with genes *CSRNP1* and *GTF2A2*).

For each trans-eGene, we ran stratified LD score regression using the top-ranked SNPs in the relevant scTBLDA factor to assess whether genetic effects were enriched in 24 functional categories [20, 22]. Although none of the enrichment p-values were significant after multiple testing correction, this was expected given only 119 individuals and a few hundred top-ranked SNPs, resulting in widespread, large standard errors (minimum BH-corrected p-value = 0.27; Fig S5). Of the 48 unique trans-eGenes, though, 15 had nominal enrichment in at least one functional annotation (enrichment estimate within the standard error range does not include zero). Notably, 11/15 (73%) of those trans-eGenes showed heritable enrichment associated with the active enhancer marker H3K27ac. This result echoes earlier findings between trans-eGenes and enhancer markers [1, 17].

To test whether expression principal components (PCs) remove broad trans-acting signals captured in our factors, we ran MatrixeQTL [23] without including expression PCs as covariates. This yielded 52 trans-eQTLs, including 34 unique trans-eGenes and 52 unique trans-eSNPs, and 35 long-range cis-eQTLs, including 35 unique cis-eGenes and 35 unique cis-eVariants (BH FDR < 0.2; Table S9). The majority of the trans-eQTLs contain a lincRNA trans-eGene (31, 60%), while the majority of the long-range cis-eQTLs follow the cis-eQTL pattern and associate with a protein-coding trans-eGene (30, 86%). We found several genes that have associations with multiple loci. For example, *LINC00936,* which inhibits the mTOR pathway (a key regulator of monocyte differentiation) [24, 25], has associations in trans with rs34217534, rs2826196, and rs10848025 in CD14+ monocytes, the first two of which are both on chromosome 21 (Bonferroni FDR ≤ 0.18). As another example, the lincRNA eGene *RP11-290F20.3,* previously associated with monocyte count and known to affect gene regulation [26], has associations in FCGR3A+ monocytes with rs555105, rs2292665, rs79552930, rs12974850, and rs348873 (Bonferroni FDR ≤ 0.18). The increase in multi-SNP associations suggests that, similar to PEER factors, including expression PCs may inadvertently remove broad-acting signal with a genuine genetic basis [**?**].

We next examined the cell-type specificity of the scTBLDA eQTLs by testing whether they replicated in a PBMC pseudo-bulk space created by averaging expression over all cells per donor. Cis-eQTLs had the highest rates of replication (188/205; 92%; BH FDR ≤ 0.2) while long-range cis- and trans-eQTLs were less likely to replicate in the pseudo-bulk space (24/42 and 37/66, respectively). Cis-eQTLs also had a higher concordance of effect direction (201/205; 98%) compared to long-range cis- and trans-eQTLs (90/108) regardless of significance. Notably, cell-type abundance was correlated with the proportion of replicated eQTLs (Spearman’s rho 0.78, p-value ≤ 0.03). The largest two cell types, CD4 T cells and CD14+ monocytes, replicated 96 and 100% of their eQTLs, respectively, in pseudobulk. This suggests that the pseudobulk analysis across all of the PBMC cell types reflects signals from the most prevalent cell types, but washes out expression patterns from less prevalent cell types (Fig 6).

For a different perspective on the cell-type specificity of eQTL associations, we tested the top features in every factor, irrespective of cell-type association, in the PBMC pseudobulk space. We found 131 cis-eQTLs with 62 unique cis-eGenes and 126 unique cis-eVariants (BH FDR < 0.2). The majority of the cis-eGenes, 123, were protein-coding while eight were lincRNA. The HLA region on chromosome 6 accounted for 30.5% of these pseudobulk PBMC cis-eQTLs. We also found four long-range cis-eQTLs that included two unique lincRNA cis-eGenes and four unique cis-eVariants, and five trans-eQTLs including five unique protein-coding trans-eGenes and five unique trans-eVariants. One of the trans-eGenes, *EIF2AK2*, has been shown to regulate transcription of SLE-associated histone and immune response genes [27]. Importantly, although some of the eQTLs had ubiquitous allelic directions and effect sizes across cell types, others did not, such as the association between *LRP10* and rs10957311 of increased expression levels with increasing minor allele count in CD14+ monocytes and pseudobulk PBMCs, but the opposite effect in B cells and FCGR3a+ monocytes. (Fig 5).

**Figure 5:**
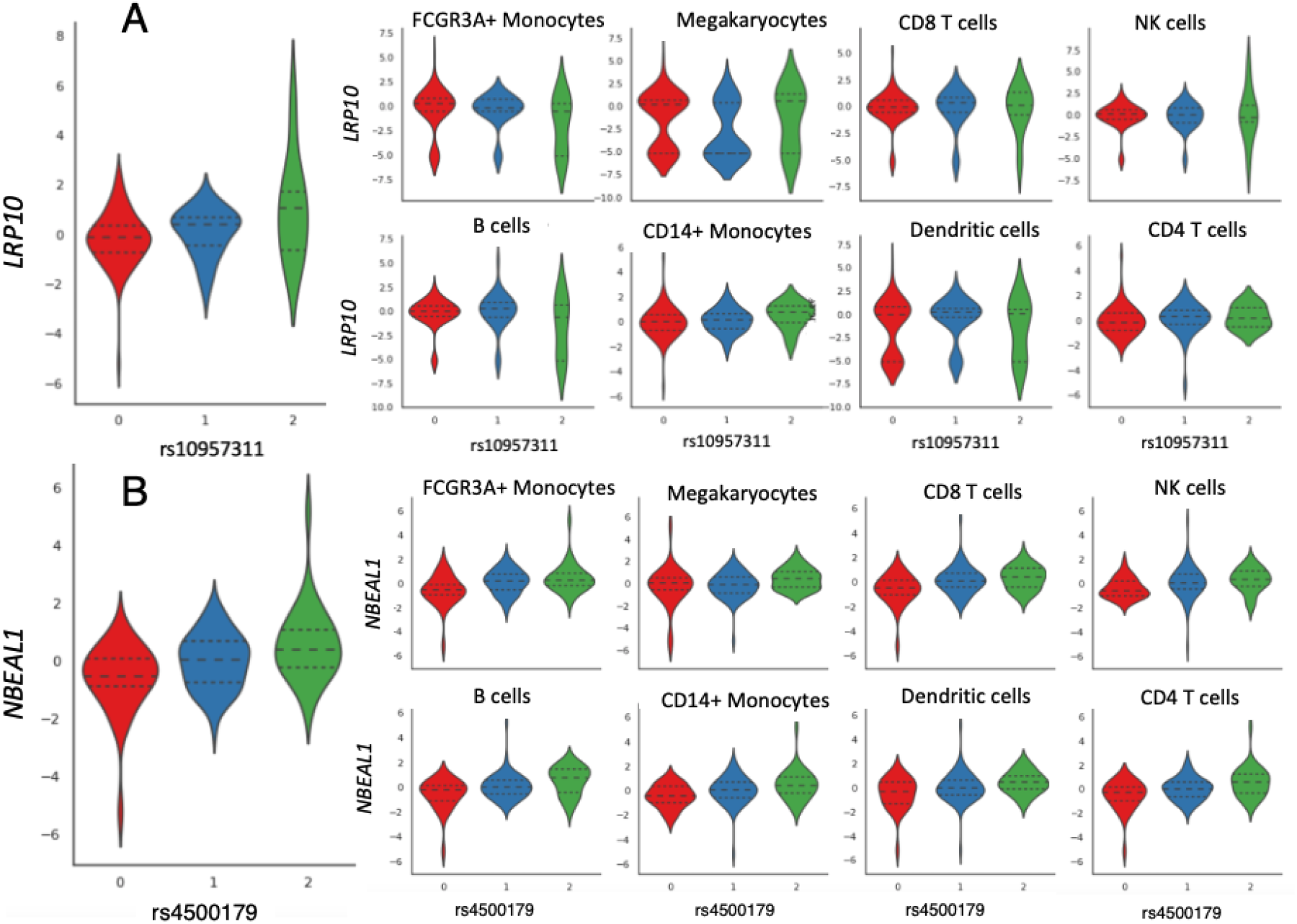
Pseudo-bulk trans-eQTLs visualized across cell types. The large plots on the left visualize the eQTLs using pseudo-bulk PBMC expression values from the testing set. The x-axis represents the number of copies of the minor allele at the given locus and the y-axis represents the quantile-normalized expression values within the given cell types for **A**. *LRP10* and **B**. *NBEAL1.*

We further explored whether our model captures cell-type proportion QTLs (ctQTLs), where the allele dosage is associated with the proportion a given cell type in a sample. We found 53 ctQTLs using the z-scored cell-type proportion as the quantitative trait (BH FDR < 0.2; Fig 6; Table S10). Natural killer (NK) and B cell proportions had the most ctQTLs, 13 and 10 respectively, while CD4 T cells only yielded one ctQTL. Chromosomes four, five, and 13 house the most ctVariants with seven, five, and five respectively. Two SNPs located ~ 10.5 kb apart on chromosome five (rs445118 and rs12655804) are both ctVariants for NK cell proportions (BH FDR ≤ 0.12). Rs445118 is a cis-eVariant for the immune complement cascade gene C6, located 2 Kb downstream, in eight tissues according to the GTEx v8 data release [28]. A ctQTL between rs811692 and CD4 T cell proportion was found in three factors (BH FDR ≤ 0.07). T cells are regulated by TGF-β signaling, which is largely mediated by *SMAD* [29]. Rs811692 is located 2 Kb upstream of *EID2,* which inhibits the formation of SMAD complexes, thereby suppressing TGF-β signaling [30]. Several ctVariants for dendritic cell proportion were located inside genes with known neuro-developmental, neuro-differentiation, and neural-related functions—*ZNF536, DSCAML1, CASQ2* [31, 32, 33].

**Figure 6:**
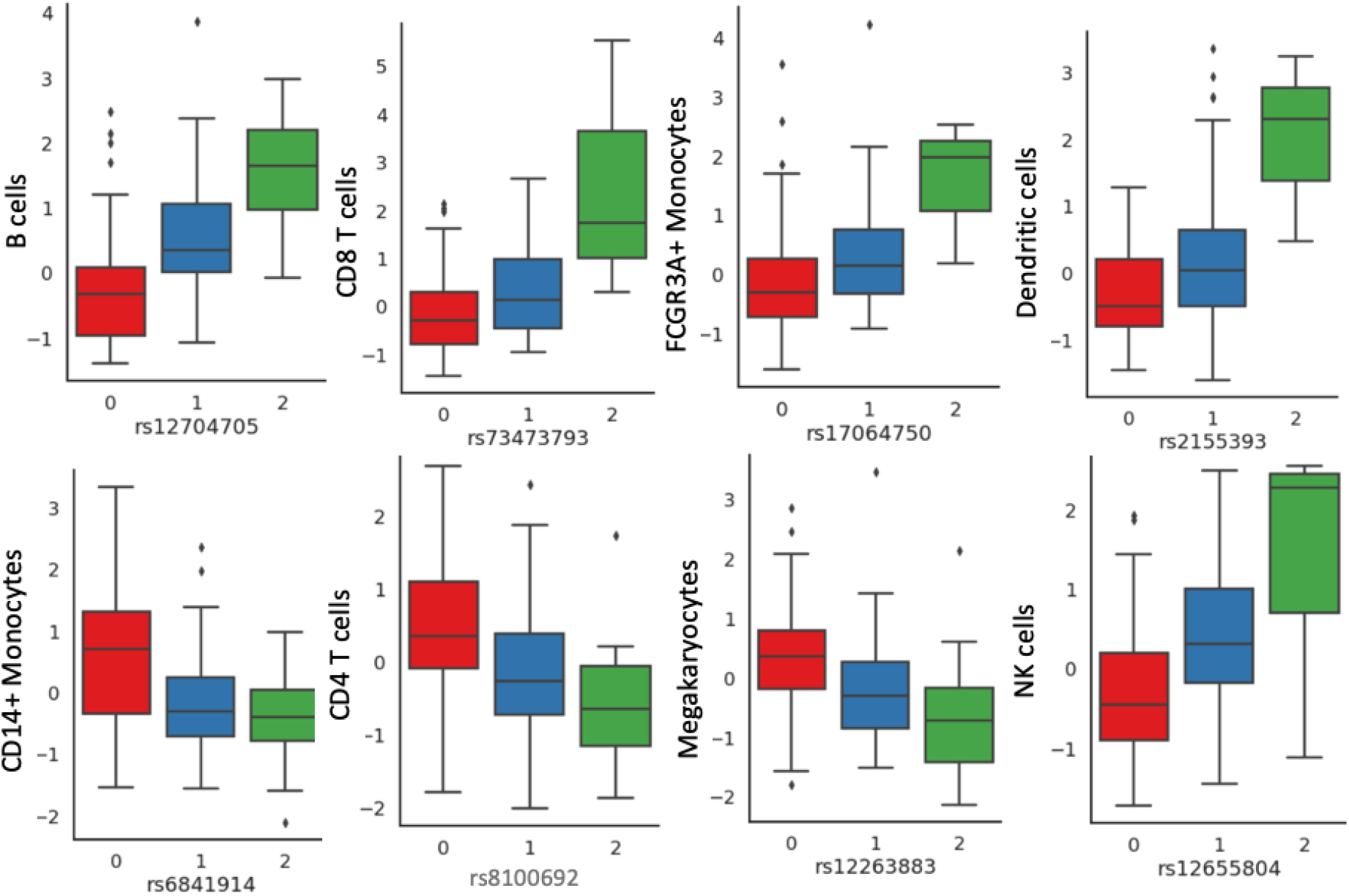
Example ctQTLs. The y-axis represents quantile normalized individual cell-type proportions for the labeled cell type. The x-axis represents the number of copies of the minor allele at the given locus.

### Replicating eQTLs in Bulk RNA-Sequencing Data

We looked for replication of the scTBLDA eQTLs in v8 GTEx bulk RNA-seq data from whole blood samples. Out of 205 TBLDA-derived cis-eQTLs, 92 (44.9%) were also identified in v8 GTEx whole blood samples. The two most abundant cell types in our single-cell data had the largest proportion of replicated cis-eQTLs, with 37/67 (55%) for CD14+ monocytes and 23/48 (48%) for CD4 T cells, suggesting that cell-type abundance affects eQTL signal in heterogeneous tissue samples. In contrast, associations in less abundant cell types may be dwarfed by conflicting or nonexistent signal from more abundant cell types (Fig 5). None of the cis-eQTLs in natural killer cells or in megakaryocytes replicate, while B cells had 15/35 (43%), dendritic cells had 9/20 (45%), CD8 T cells had 6/15 (0.4%), and FCGR3A+ monocytes had 2/9 (22%) replicate in the bulk RNA-seq GTEx whole blood samples.

High proportions of scTBLDA eGenes and eVariants were present in the GTEx whole blood eQTL list; 77/90 (85.5%) cis-eGenes and 116/169 (68.6%) cis-eVariants were part of a GTEx cis-eQTL in whole blood. Although the 77 overlapping scTBLDA cis-eGenes account for only 0.62% of the GTEx whole blood cis-eGenes, they are disproportionately part of 2.9% of all cis-eQTLs. Similarly, the 116 cis-eVariants only make up 9.1 × 10^-3^% of all unique GTEx cis-eVariants, but are over represented as 0.023% of all unique GTEx cis-eQTLs. None of the scTBLDA trans-eQTLs reach significance in the genome-wide GTEx whole blood trans-eQTL list (which only has 140 trans-eQTLs). Of the five scTBLDA PBMC pseudobulk trans-eQTLs, none replicated in the GTEx whole blood eQTLs. This may be due to differing cell-type populations and cohort ancestry or because these trans-eQTLs are specific to SLE patients.

We then examined bulk RNA-seq expression data from GTEx EBV-transformed lymphocytes (LCLs), which is a cultured line of a single-cell type. Of the 205 scTBLDA-derived cis-eQTLs, 39 (19%) were also identified by the GTEx consortium in LCL bulk-RNAseq data. Mirroring the whole blood results, the scTBLDA cis-eGenes and cis-eVariants were enriched among the GTEx cis-eQTLs. While the 41 overlapping cis-eGenes account for merely 0.83% of the unique GTEx LCL cis-eGenes, 4.4 % of LCL cis-eQTLs contained one of those eGenes. Similarly, the 58 cis-eVariants make up 0.019% of the unique e-Variants but 0.03% of all cis-eVariants. Because lymphocytes are B cells, we expect B cell eQTLs to have the highest overlap. Although 10 (28.6%) of the TBLDA B cell cis-eQTLs were present in the GTEx LCL list, dendritic cells actually had the highest proportion of cis-eQTL replication—40%—although the net number was still low (eight). Over half (3/5) of testable scTBLDA B cell trans-eQTLs replicate in the GTEx LCL data (BH FDR ≤ 0.2), demonstrating higher replicability of cell-type specific versus pseudobulk trans-eQTLs.

Next, we checked if the scTBLDA eQTLs were found in three prior PBMC eQTL studies. The Vosa study mapped cis- and trans-eQTLs in whole blood from 31,684 individuals [34]. Of the unique scTBLDA trans-eGenes, 21/48 (44%) were trans-eGenes in the Vosa study, 17/39 (44%) of the long-range cis-eGenes were trans-eGenes in th Vosa study, and 47/90 (52%) of unique scTBLDA cis-eGenes had trans-eQTLs in the Vosa study. The TBLDA eGenes are over-represented in the Vosa study; they comprise 1.2% of the unique eGenes, but 1.9% of the eGenes overall. In particular, the TBLDA trans-eGenes are represented in the Vosa study double versus what was expected— they make up 0.3% of the Vosa study unique eGenes but 0.7% of the eQTLs. Three of the scTBLDA cis-eVariants were trans-eVariants here, but no TBLDA trans-eVariants were eVariants in the Vosa study. However, the Vosa study only tested a subset of 10K trait-associated SNPs as trans-eVariants, only 391 of which overlapped with our LD-pruned SNP set. It should also be noted that only 0.03% (37/115473) of their whole blood trans-eQTLs had significant replication (FDR ≤ 0.05) in scRNA-seq data sets.

The Van Der Wijst study conducted cis-eQTL association mapping in 25K PBMCs from 45 individuals [10]. Again, we found overlapping and enriched signal–29/292 (10%) of the unique cis-eGenes in the van Der Wijst study were scTBLDA eGenes, and they accounted for 28.6% of the cis-eQTLs. Interestingly, while more scTBLDA cell-type specific trans-eGenes were found as cis-eGenes here than scTBLDA pseudobulk PBMC trans-eGenes (six versus zero), more scTBLDA PBMC cis-eQTLs overlapped with these cis-eGenes than scTBLDA cell-type specific cis-eGenes (26 versus 11). Indeed, the van Der Wijst study found that 87.3% of their cis-eQTLs were also significant in their pseudobulk PBMC-based association mapping. We both found a shared region on chromosome 4 (118977976 to 119219187) containing several cis-associations with *SNHG8* in CD4 T cells (four in the van Der Wijst study, seven scTBLDA eQTLs).

The Fairfax study looked at mapped cis- and trans-eQTLs in purified monocytes and B cells from 288 healthy European individuals [35]. Although none of our trans-eQTLs were identified in the study, the trans-eGenes that were shared, *AOAH* and *NCF2,* were part of a disproportionate percentage of the Fairfax study’s trans-eQTLs—they comprised 1.03% (2/195) of the unique trans-eGenes but 5.4% (92/1704) of trans-eQTLs overall. Several identified eGenes and genomic regions overlapped between our results, suggesting common regulatory networks. For example, we both found a region on chromosome 8, between 145875231 and 146128143, that had multiple cis-associations with *RPL8* in CD14+ monocytes and B cells. Despite our data set containing eight cell types, scTBLDA identified these interactions only in CD14+ monocytes, B cells, and dendritic cells, which are closest to monocytes. This cis association was also present in the van Der Wijst study [10], although it did not reach significance in monocytes or dendritic cells despite having low p-values (p ≤ 0.001, 0.01 respectively), demonstrating increased power in our analysis to detect associations that are indeed present across data sets and cell types. scTBLDA further identified a trans-eQTL between *RPL8* and rs73147705. Another shared eGene, *ITGB2*, which encodes an integrin chain and is associated with leukocyte adhesion deficiency, only had cis-associations in the Fairfax study. scTBLDA not only identified a cis-eQTL but also found a trans association in natural killer cells with rs2864754 (Bonferroni FDR ≤ 0.054). Taken together, these results suggest that the eQTL results from scTBLDA broadly replicate across other whole blood eQTL studies while expanding their findings.

## Discussion

In this paper, we extend our prior model, TBLDA, to the various practical aspects of scRNA-seq data in order to create a method that allows scRNA-seq count data and genotype information to be used for eQTL mapping. We adapt our method to use GPUs to speed up our approach by two orders of magnitude. We include sparsity-inducing priors in the factorization to account for the large number of zero reads per gene across cells and to aid in the interpretation of the results. We incorporate amortization into the variational approximation in order to avoid learning *N* local variational parameters—essential when data containing millions of cells are increasingly common. The resulting method, scTBLDA, can fit expression from nearly a half million cells and tens of thousands of SNPs within 35 minutes when run on a GPU. To the best of our knowledge, this is the first exploration of the relationships between genotype and single-cell expression data that does not use pseudobulk-based association mapping.

We fit our scTBLDA model to nearly 400K PBMCs and 154K SNPs from 119 SLE patients from a previous paper [18], finding 1320 cell-type associated factors. SNPs and genes that appear across multiple cell-type associated factors were enriched for immune-related and regulatory gene sets as well as local genomic regions and various SNP classes, notably enhancer-related classes. We demonstrated that scTBLDA factors can identify genetic variants associated with gene expression levels unique to subtypes within labeled cell types, including an association between the lncRNA *SNHG8* and rs7241781 in a subset of B cells. Using the top SNPs and genes loaded onto each factor, we mapped 205 cis-eQTLs, 42 long-range cis-eQTLs, and 66 trans-eQTLs. Although nearly all (98%) of the cis-eQTLs replicated in pseudo-bulk PBMC data, only half (56%) of our trans-eQTLs did. This highlights the importance of looking at associations within specific cell types instead of the heterogeneous mixture given by bulk RNA-seq data. Further, we discovered 53 cell-type proportion QTLs, including a fascinating link between rs811692 and CD4 T cell proportion that may involve SMAD-mediated TFG-β signaling.

The scTBLDA-discovered eGenes and eVariants were enriched across four other eQTL studies comprising both bulk and scRNA-seq data [1, 35, 34, 10]. We replicate our trans-eQTLs in the GTEx whole blood and LCL eQTL data; in particular, three of our five scTBLDA B cell trans-eQTLs replicate in the v8 GTEx LCL data. There are many potential reasons for the lack of complete eQTL overlap between studies, including minimal overlap in starting variant sets, different population ancestries, modified regulation in healthy versus SLE cohorts, and statistical power differences. We suspect that there are additional eQTLs in our results that could not be identified because the held-out data used for discovery was underpowered, the cell-type labels were imprecise, or because the number of features did not allow the comprehensive capture of every eQTL across the sample. Despite these false negatives, our model opens the door to an orthogonal approach to eQTL discovery that will require methods and strategies for discovery, validation, and replication that differ from the current paradigm for eQTL discovery. We have merely scratched the surface in this paper.

Our approach has drawbacks and assumptions that should be mentioned. One of the largest is that the specificity of the factors is bounded by their number. There must be a balance between the number of factors being too large, creating factors defined by too few genes to be meaningful, and too small, overloading the biological signal in each factor to include multiple biological processes. Rather than comprehensively finding all possible one-to-one associations, scTBLDA jointly decomposes both modalities to identify groups of associated features. These analyses can be useful for exploratory data analysis such that the factors become a starting point to investigate potential regulatory mechanisms. Another limitation in scTBLDA lies in the eQTL discovery phase, and in particular in taking the given cell-type labels as ground truth. Our results show that some cells appear to have factor proportions most characteristic of another cell type (Fig 1), and that some cell types have clear subgroup structures that confound discovery in our eQTL pipeline. By not taking cell-type label uncertainty into account, the signal for downstream eQTL mapping may be adulterated by including mislabeled cells in the cell-type specific expression statistics. While we mapped eQTLs in factors enriched for single cell types based on factor-cell type associations, our process does not preclude general eQTLs. Further, our validation still relies on univariate tests that explicitly control for ancestry, large effect confounders, and other sources of noise. The approaches used to control for these possible confounders may remove actual signal that scTBLDA is able to identify in the factors but we are not able to harness for discovery using the classic univariate mapping pipeline. Finally, validating the bimodal associations using one-to-one eQTL mapping does not directly validate the biological signal found in multi-gene, multi-variant factors, which are the heart of this method. We expect many single-feature associations to fall below significance thresholds given only 119 individuals and the small held-out expression test set used for eQTL mapping.

Many heritable diseases are thought to arise based on expression modifications from large numbers of small trans effects, each of which individually would not pass significance testing even with a huge sample size [36]. Applied to healthy and diseased cohorts, scTBLDA can provide an alternative framework to univariate association testing approaches to study the genetic regulation of complex traits through many trans effects. Further, scTBLDA may be applied to any count-based, multimodal nested data; in the current framework it is simple to swap out the SNP-inspired binomial likelihood to fit another distribution. Going forward, scTBLDA can be applied to a wide variety of data sets and inspire wet-lab experiments to validate proposed associations.

## Extended Methods

### Data QC and Feature Selection

We excluded all cells with over 10% of transcripts coming from mitochondrial genes, as high proportions of mitochondrial gene reads indicate low-quality or dying cells. Cells with more than 12,500 total UMIs were also discarded. This resulted in 400,280 cells for downstream analysis.

Next, we considered all autosomal protein-coding and lncRNA genes that were present in at least 0.25% (1000) of the cells. We ranked the genes according to the deviancePoisson function [37] and selected the top 2500. We randomly selected max(5, 15%(number of cells from individual *i* of cell type z)) cells from each individual of each cell type to hold out for the eQTL pipeline.

We pruned the SNPs on each chromosome using plink v1.9 [38], running `-indep-pairwise with` *r*^2^ = 0.2, *bp_window_size* = 200(*Kb*), and *stepsize* = 10. This resulted in a final set of 154,126 LD-pruned SNPs.

### Ancestry Structure

We ran terastructure [39] on the 3,725,027 SNPs with the following options: `-rfreq=745000 and -K=2` to produce the genotype-specific portion of the model in the same way as in earlier work [17].

### scTBLDA

While the generative model is the same as TBLDA, the variational approximate distribution for scTBLDA is specified in a mean-field approach as follows:

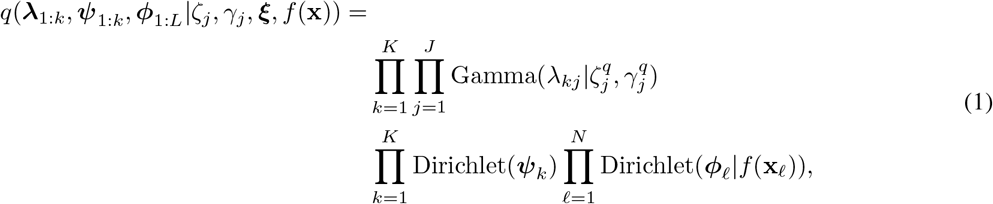

where *J* is the total number of SNPs, *λ_jk_* ~ Beta(*ζ_j_*, *γ_j_*) is the weight for SNP *j* in topic *k*, and *ψ_k_* ~ Dirichlet(*ξ*) is the *k*th gene expression factor.

### Model Runs

We used Pyro v1.4.0 [40]’s stochastic variational inference framework to fit the model, using `pyro.poutine.scale` (scale=1.0 × *L* * *G*), where *L* is the number of cells and *G* is the number of genes, for numerical stability, a clipped Adam optimizer, 200 minibatches, a learning rate of 2 × 10^-4^, *σ* = 1, *δ* = 0.05, and *μ* = 0.5. The model was fit separately for SNP sets from each chromosome, for a total of 22 runs. Code to run scTBLDA is available at https://github.com/gewirtz/scTBLDA.

### Feature Ranking

For each factor and modality, we ran quantile regression using `statsmodels.formula.api.quantreg` on MAF or average expression level at the 0.9th quantile. SNPs with loadings greater than expected from the regression line were considered top features. Genes with loadings greater than the regression line and with log(loadings) > −10.2 were considered top features (Fig. S2). In cases where fewer than 10% of genes were selected and additional genes were available above the −10.2 threshold, the quantile was reduced by 0.05 and the regression was repeated.

### Covariate-Factor Association

We computed the inner product between an indicator matrix depicting cell membership in each covariate and the estimated cell-factor proportions, Φ, using the encoder. The inner product was normalized such that each factor’s values lay on the simplex (over covariates). For batch, we calculated the full log probability for each factor given a Dirichlet distribution with parameter *α_i_* equal to the number of cells labeled with covariate *i*. All factors with log probability < −60000 were considered associated with batch and removed (Fig. S3). For cell type, we calculated the unnormalized log probability for each factor and each cell type given a Dirichlet distribution with *α_i_* as the log of the number of cells assigned to covariate *i*. The largest 15% of the log probabilities were considered cell-type associated factors.

Cell-type associated features are all genes and SNPs that were top-ranked in all or over 75%, respectively, of each cell type’s associated factors.

### QTL Mapping

We used MatrixEQTL v2.3 [23] `with modelLINEAR` to run the eQTL testing. Size factors for each cell were computed using the scran [41] function `calculateSumFactors` with clusters set to the labeled cell types. The average log(1 + adjusted count) was computed for all cells of each cell type within individuals, and expression for all genes was quantile-normalized to a standard Gaussian. Following earlier work [42], the first 15 expression PCs were included as covariates, along with the first genotype PC after scaling the genotype matrix [43]. FDR was computed using the Benjamini-Hochberg procedure over tests in each run for cis- and trans-eQTL and protein-coding and lncRNA genes separately. For ctQTLs, we used the first genotype PC as a covariate and tested the top SNPs in cell-type associated factors against the z-scored cell-type proportions for each individual. Where necessary to compare to prior studies, we used pyliftover version 0.4 to convert between GChr37 and GChr38.

## Supporting information

**Figure S1:**
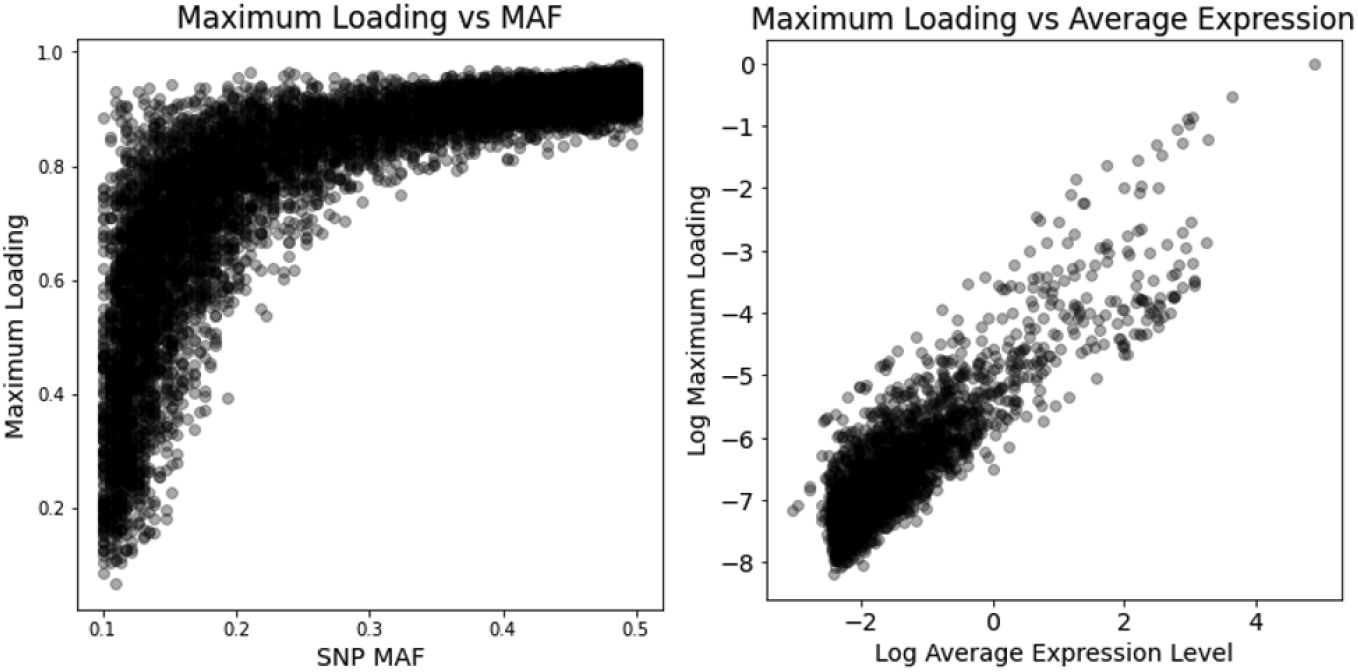
Relationship between maximum loading weight and feature frequency. Feature frequency is on the x-axis and the y-axis represents the maximum loading for each feature across all factors.

**Table S1:** Evaluation of activation functions in the encoder. Runs were done with learning rate of 0.0005, two hidden layers with 1000 and 200 neurons, and SNPs from chromosome one. Model loss on the held-out cells and root mean squared error on the posterior expression matrix was lowest for sigmoid. All tested activation functions produced similar loss values, though ReLU had a notably higher expression RMSE.

| Activation Function | Test Set Loss | Test Expression RMSE |
| --- | --- | --- |
| Sigmoid | 0.0661 | 0.2926 |
| Softplus | 0.0667 | 0.3138 |
| ReLU | 0.0702 | 0.5967 |

**Table S2:** Evaluation of encoder hidden layer sizes. Model loss on the held-out cells and root mean squared error on the posterior expression matrix are recorded for runs at learning rate = 0.0005 using sigmoid activation and SNPs from chromosome one.

| Hidden Layer Sizes | Test Set Loss | Test Expression RMSE |
| --- | --- | --- |
| 150 - 150 | 0.0662 | 0.2796 |
| 300 - 150 | 0.0661 | 0.2759 |
| 500 - 200 | 0.0660 | 0.3282 |
| 1000 - 200 | 0.0661 | 0.2926 |
| 1000 - 500 | 0.0661 | 0.2960 |

**Table S3:** Evaluation of varied learning rates. Model loss on the held-out cells and root mean squared error on the posterior expression matrix are recorded for runs using sigmoid activation, hidden layers of 300 and 150 neurons, and SNPs from chromosome one.

| Learning Rate | Test Set Loss | Test Expression RMSE |
| --- | --- | --- |
| 0.0001 | 0.0662 | 0.2826 |
| 0.00025 | 0.0659 | 0.2806 |
| 0.0005 | 0.0661 | 0.2759 |
| 0.001 | 0.0660 | 0.2800 |
| 0.002 | 0.0660 | 0.2790 |

**Figure S2:**
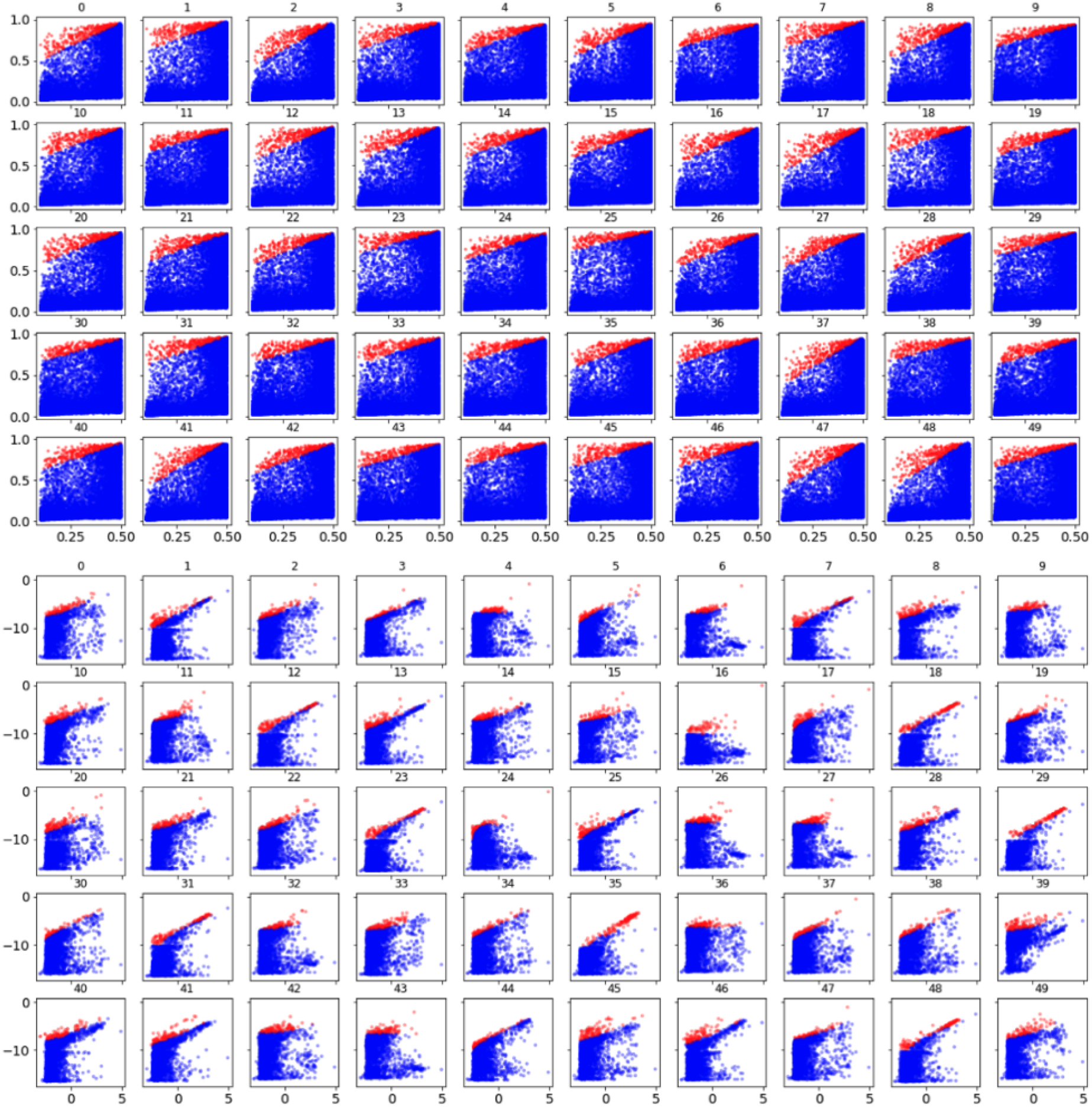
Ranking features per factor using quantile regression. Top: Each plot depicts loading on the y-axis and MAF on the x-axis for a given factor, labeled as the subplot title, for the run using SNPs from chromosome one. Bottom: Each plot depicts log(loading) on the y-axis and log(average expression level) on the x-axis for a given factor, labeled as the subplot title, for the run using SNPs from chromosome one. Genes and SNPs chosen as top features for each factor are colored red.

**Table S4:** Evaluation of adding dropout to the encoder. Model loss on the held-out cells and root mean squared error on the posterior expression matrix are recorded for runs at learning rate = 0.0005 using sigmoid activation, hidden layers of 300 and 150 neurons, and SNPs from chromosome one.

| Dropout Proportion | Test Set Loss | Test Expression RMSE |
| --- | --- | --- |
| 0 | 0.0661 | 0.2759 |
| 0.1 | 0.0663 | 0.2836 |
| 0.2 | 0.0664 | 0.2924 |
| 0.3 | 0.0669 | 0.2996 |

**Figure S3:**
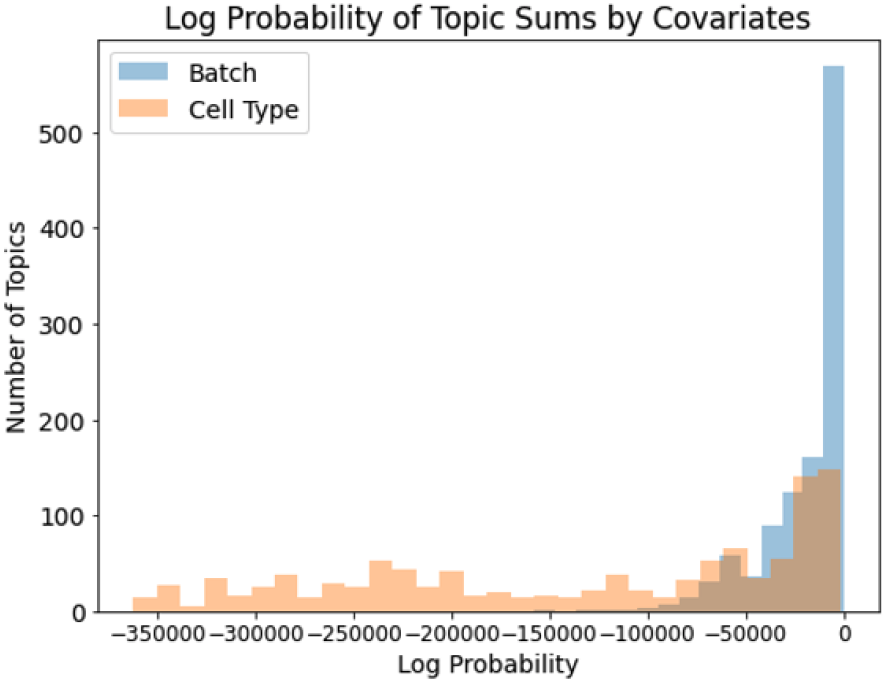
Histogram depicting the summed log probability of batch and cell type for factors across all runs.

**Figure S4:**
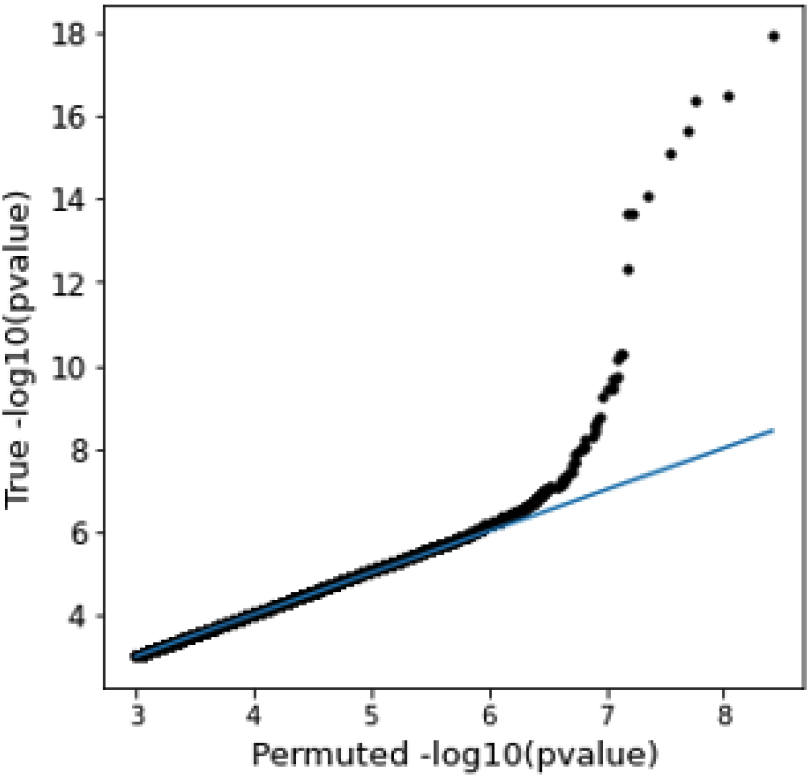
Enriched p-values between top features on the same factors. Enrichment in low Matrix eQTL p-values for trans- and cis-eQTLs using the top features on common factors compared with permuting the expression values and covariates.

**Figure S5:**
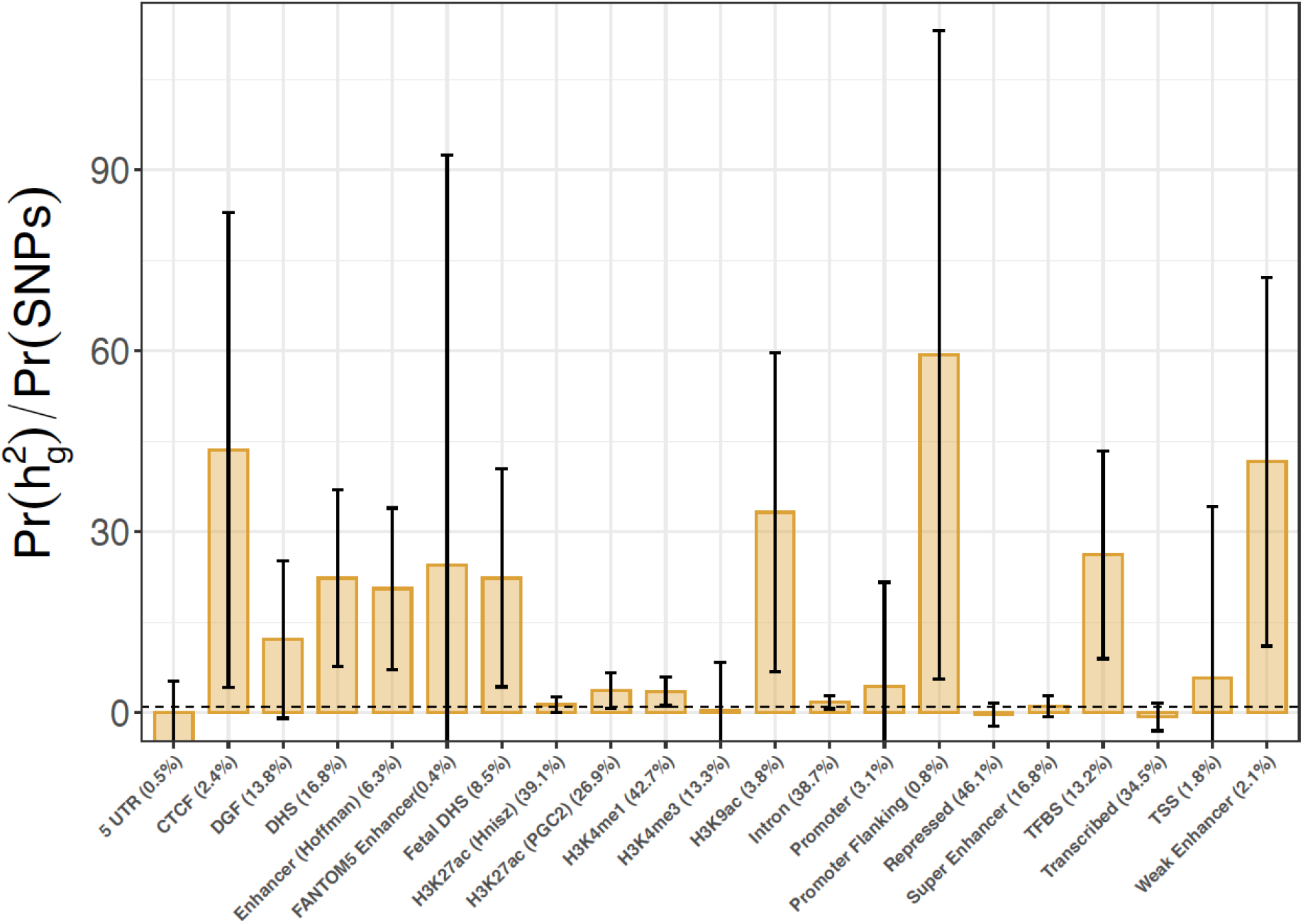
Stratified LD-score regression enrichment analysis on 24 functional categories using the top-ranked SNPs for the trans-eGene *LINC00861* in CD4 T cells. The proportion of SNPs colocalizing with each annotation category is listed in parentheses. Categories with standard errors below 0 were removed prior to plotting. The minimum BH-corrected p-value (0.27) across all trans-eGenes came from the proportion of heritability in *LINC00861* expression levels explained by top-ranked SNPs in TFBS.

**Figure S6:**
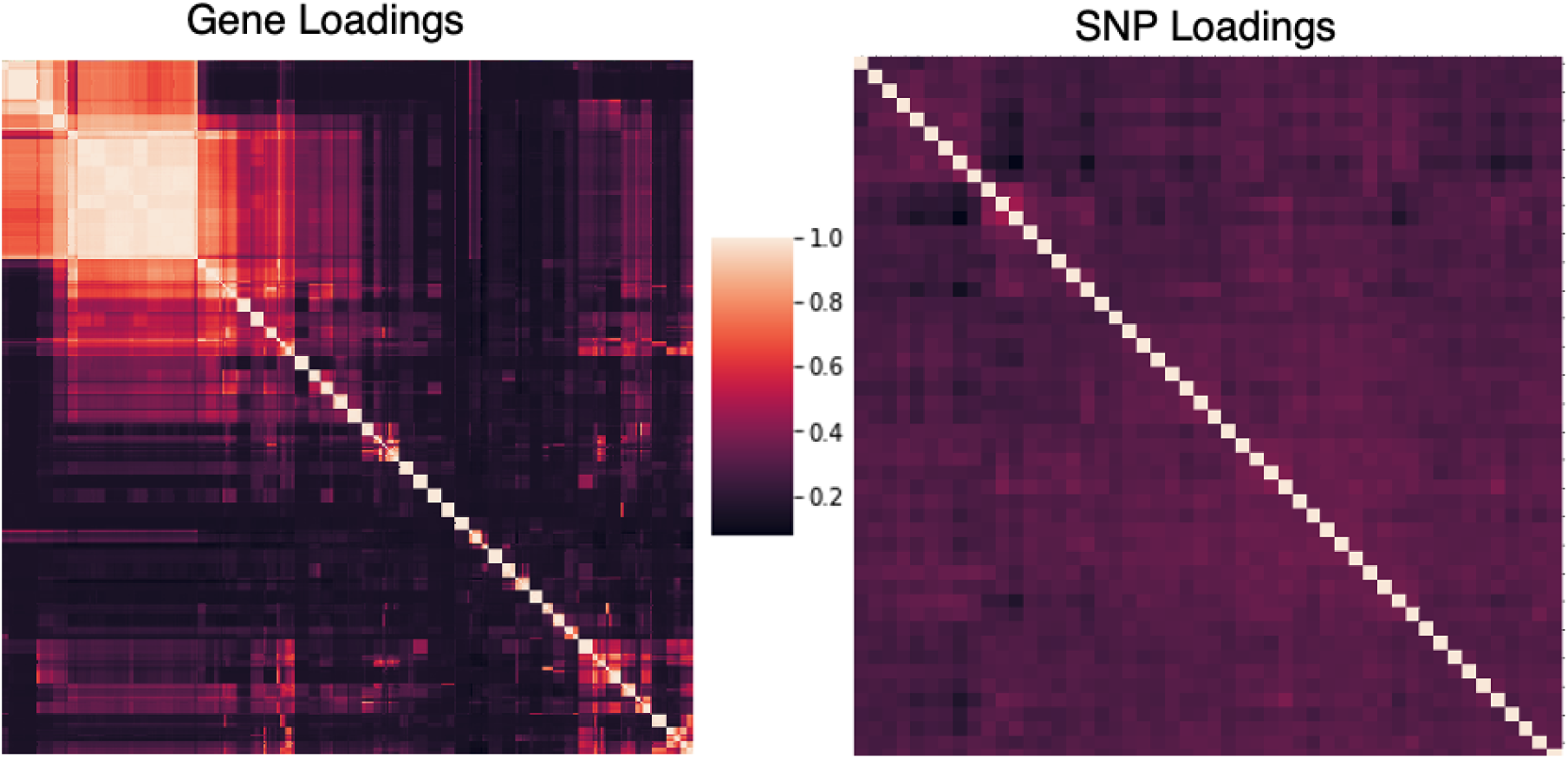
Correlations between gene and SNP loadings. Left: Heatmap depicting the Pearson correlation among gene loadings for factors across all scTBLDA runs. Right: Heatmap depicting the Pearson correlation among chromosome 12 SNP loadings for learned factors. factors are hierarchically-clustered. The present correlation structure justifies looking for enrichment by fully permuting expression levels.

## Acknowledgments

AG, FWT, and BEE were funded by Helmsley Trust grant AWD1006624, NIH NCI 5U2CCA233195, NIH NHLBI R01 HL133218, and NSF CAREER AWD1005627.

